# Rigorous anterograde trans-monosynaptic tracing of genetic defined neurons with retargeted HSV1 H129

**DOI:** 10.1101/2020.12.01.407312

**Authors:** Peng Su, Min Ying, Jinjin Xia, Yingli Li, Yang Wu, Huadong Wang, Fuqiang Xu

## Abstract

Neuroanatomical tracing technology is fundamental for unraveling the complex network of brain connectome. Tracing tools that could spread between neurons are urgently needed, especially the rigorous trans-monosynaptic anterograde tracer is still lacking. HSV1 strain H129 was proved to be an anterograde tracer and has been used to trace neuronal networks in several reports. However, H129 has a serious defect that it was demonstrated to infect neurons via axon terminals. Thus, when using H129 to dissect output neural circuit, its terminal take up capacity should be carefully considered. Here, we report a recombinant H129 that carrying the anti-Her2 scFv in glycoprotein D to target genetically defined neurons. With the usage of helper virus complementarily expressing Her2 and gD, we can realize the elucidation of direct projection regions of either a given brain nucleus or a specific neuron type. The retargeted H129 system complements the current neural circuit tracer arsenal, which provides a rigorous and practical anterograde trans-monosynaptic tool.

## Introduction

Transneuronal tracing of neural circuit using neuroinvasive viral tools is gradually becoming a powerful approach for defining the synaptic organization of neural pathways ^1, 2^. The method utilize the ability of viruses to invade neurons and produce infectious progenies that pass transneuronally at synapses to infect synaptically connected neurons. As the virus replicates and spreads through the neural networks, neurons are labeled by viral or reporter proteins, thus making the definition of synaptic organization of functionally defined neural circuitry possible.

Mapping the neuronal circuits requires both anterograde and retrograde tracers. So far, rabies virus (RV) ^3^ and pseudorabies virus (PRV) ^4^ have been modified to be used as retrograde mono-synaptic and retrograde multi-synaptic tracers, respectively. Meanwhile, vesicular stomatitis virus (VSV) ^5^ and H129 ^6^, an HSV1 isolate from the brain of an individual who suffered from HSV1 viral encephalitis, were reported to be anterograde polysynaptic tracers. Kevin T. Beier et al. proved that VSV performed an anterograde transmission manner and has been used to trace neuronal networks in several reports ^7, 8^. Liching Lo and David J. Anderson introduced a H129 mutant named H129ΔTK-TT which could map output networks started from genetic defined neurons ^9^. Recently, a H129 mutant, H129-ΔTK, has been applied as an anterograde monosynaptic tracer ^10^. However, H129 has a serious defect that HSV1 was demonstrated to invade neurons via axon terminals ^11, 12^. Thus, when using H129 to dissect neuronal circuit, its terminal take up capacity should be carefully considered. Eliminating the terminal take up capacity of H129 is an important solution for its better application, but how? We may find answer in oncolytic HSV (oHSV) researches. The first-generation retargeted oHSVs carry the retargeting ligand in gD, in place of a portion of the glycoprotein ^13, 14^. The deleted sequence is critical for gD interaction with its natural receptors HVEM (herpesvirus entry mediator) and nectin1. The resulting recombinants are detargeted from these receptors. Gabrielle and Bernard have reported a series of researches of retargeted oHSV based on chimeric glycoprotein D, mainly via the insertion of a scFv (single chain antibody variable region) in gDΔ6–38 ^15^. Thus, we could possibly use a retargeted H129 to target genetic defined neurons. According to previous reports, we chosed human Her2 (Human epidermal growth factor receptor 2) protein, which was widely studied and barely expressed in brain, as the target of retargeted H129. Using the recombinants carrying the anti-Her2 scFv in engineered gD protein and helper virus complementarily expressing Her2 and gD, we can realize the dissection of direct projection targets of either a given brain nucleus or a specific neuron type.

## Results

### Construction of gD null and Her2 targeting recombinant H129

In order to facilitate the subsequent construction of Her2 targeting H129 virus, we first constructed a gD null H129 strain H129ΔgD-tdTomato, and named it Hs01. Hs01 was generated by replacing the gD gene of H129 with the red fluorescent protein tdTomato (tdT) expressing box (Fig.1A). Firstly, we constructed BHK-gD cell line which expressed gD stably. Next, the shuttle vector pHs01 and H129 virus were recombined in BHK-gD cell, and the recombinant Hs01 expressing red fluorescence was purified (Fig.1B). In the process of plaque purification, due to the faster replication rate of H129, it should be more patient and more accurate in picking the plaques of Hs01. In order to test the infection properties, Hs01 was used to infect BHk-gD, primary neurons and BHK cells, respectively. The results showed that Hs01 could only proliferate in BHK-gD cells. Though Hs01 could infect primary neurons and BHK cells, it could not produce infectious progeny virus again (Fig.1C). Therefore, by infecting BHK cells, we could test whether the acquired Hs01 virus was a fully purified monoclonal strain.

**Figure 1.**
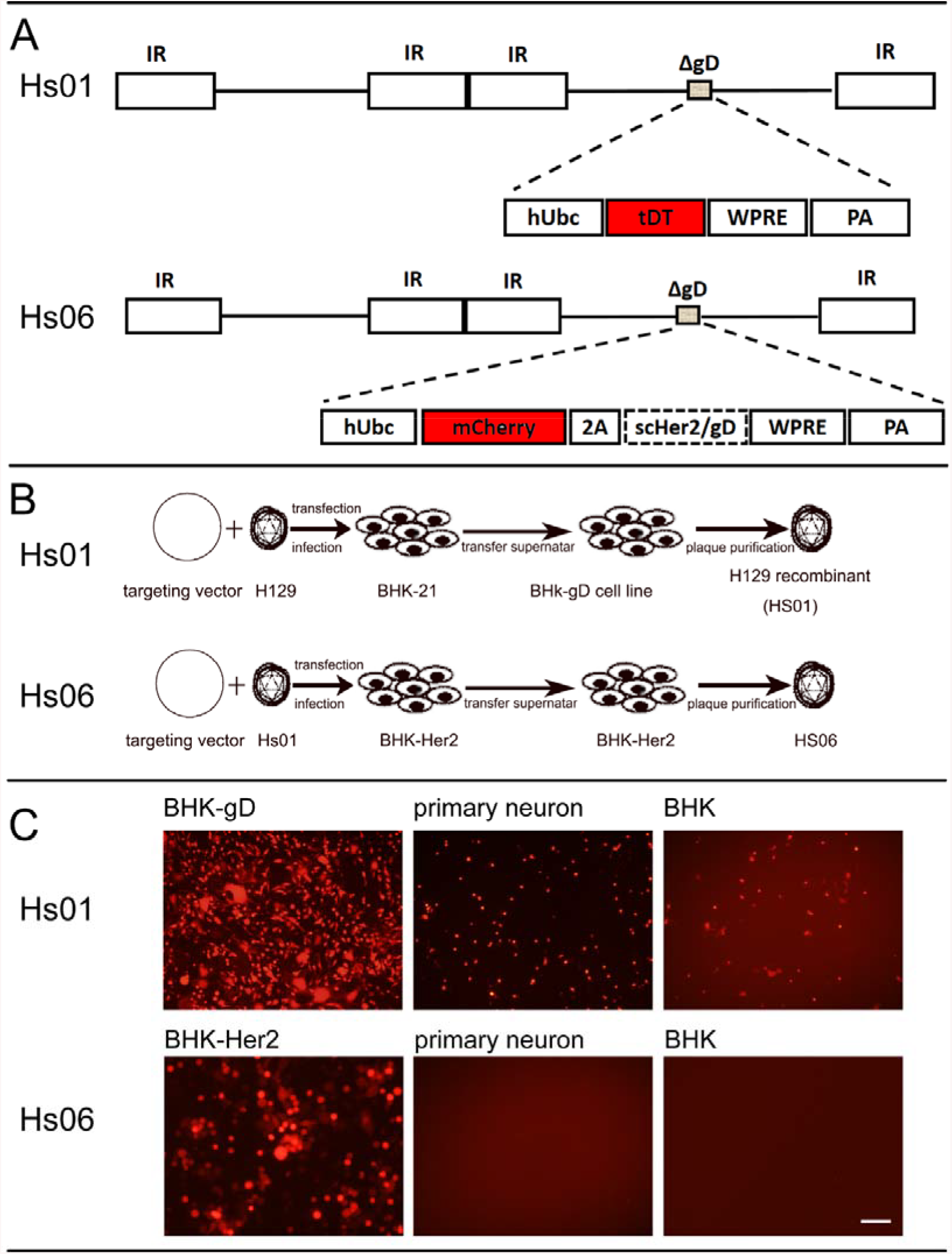
Construction and identification of gD null and Her2 targeting H129 recombinant. (A) Schematic diagrams of Hs01 and Hs06 genome. (B) Hs01 and Hs06 were generated by recombination and plaque purification. pHs01 and H129 recombine in the BHK-gD cells, then plaque purification was performed to gain strain Hs01. pHs06 and Hs01 recombine in the BHK-Her2 cells, then plaque purification was performed to gain strain Hs06. (C) Cell tropism of Hs01 and Hs06. Hs01 and Hs06 infected BHK-gD, primary neurons and BHK cells, respectively. Hs01 could replicate in BHK-gD cells and could only infect once in primary neurons or BHK cells. Hs06 could replicate in BHK-Her2 cells and could not infect primary neurons or BHK cells. Scale bar, 100 μm.

To construct the recombinant H129 specifically targeting Her2, we first constructed shuttle vector pHs06 (Fig.1A) and cell line BHK-Her2 which stably expressing Her2 protein. Hs01 and pHs06 were recombined in BHK-Her2 cells to generate strain Hs06 (Fig.1B). The obtained Hs06 was used to infect BHK-Her2 cells, primary neurons and BHK cells, respectively. The results showed that Hs06 could only infect BHK-Her2 cells, indicating that Hs06 had the specificity of targeting the Her2 receptor (Fig.1C). Since recombinant H129 strain Hs06 could only recognize cells that express Her2 receptor, and the neurons of animal central nervous system hardly expressed Her2 receptor, we had achieved the ability to eliminate the axon terminal infection of H129.

### Shortened Her2 receptor could be recognized by Hs06

At present, the virus commonly used for exogenous gene expression in the central nervous system is adeno-associated virus (AAV) ^16^. However, its packaging capacity is small, and it can only contain the exogenous gene expression box of no more than 5 Kb. The complete Her2 receptor, however, consisted of 1,255 amino acids and has a gene length of 3,768 bp. Since the gene expression box also contains other sequences such as promoters and terminator sequence, Her2 gene is difficult to be carried by AAV vector. If small promoters were used, the expressing level would be low and we could not add a fluorescent protein gene to indicate the expression of Her2 receptor. So could we shorten the length of Her2 receptor? By analyzing the protein structure of Her2, we found it had a long intracellular region of 580 amino acids. Hs06 only needed to recognize the extracellular region of Her2 to adsorb to the cell surface and initiate subsequent infection process. Therefore, we predicted that the ability of Her2-mediated infection of Hs06 would not be affected after removal of the intracellular region of Her2. For the construction of the shortened Her2, 9 amino acids of the intracellular region were retained to avoid changes in the structure of the transmembrane region (Fig.2A). The shortened Her2 receptor was named Her2CT9. Since the length of Her2CT9 is reduced to 2,055 bp, we could easily add fluorescent protein gene when constructing its AAV expression vector. After obtaining Her2CT9, we constructed the cell line BHK-Her2CT9 for subsequent verification experiments. Hs06 was used to infect cells expressing Her2 or Her2CT9, respectively, and the results showed that the modified Her2CT9 receptor could effectively mediate the entry of Hs06 into cells (Fig.2B).

**Figure 2.**
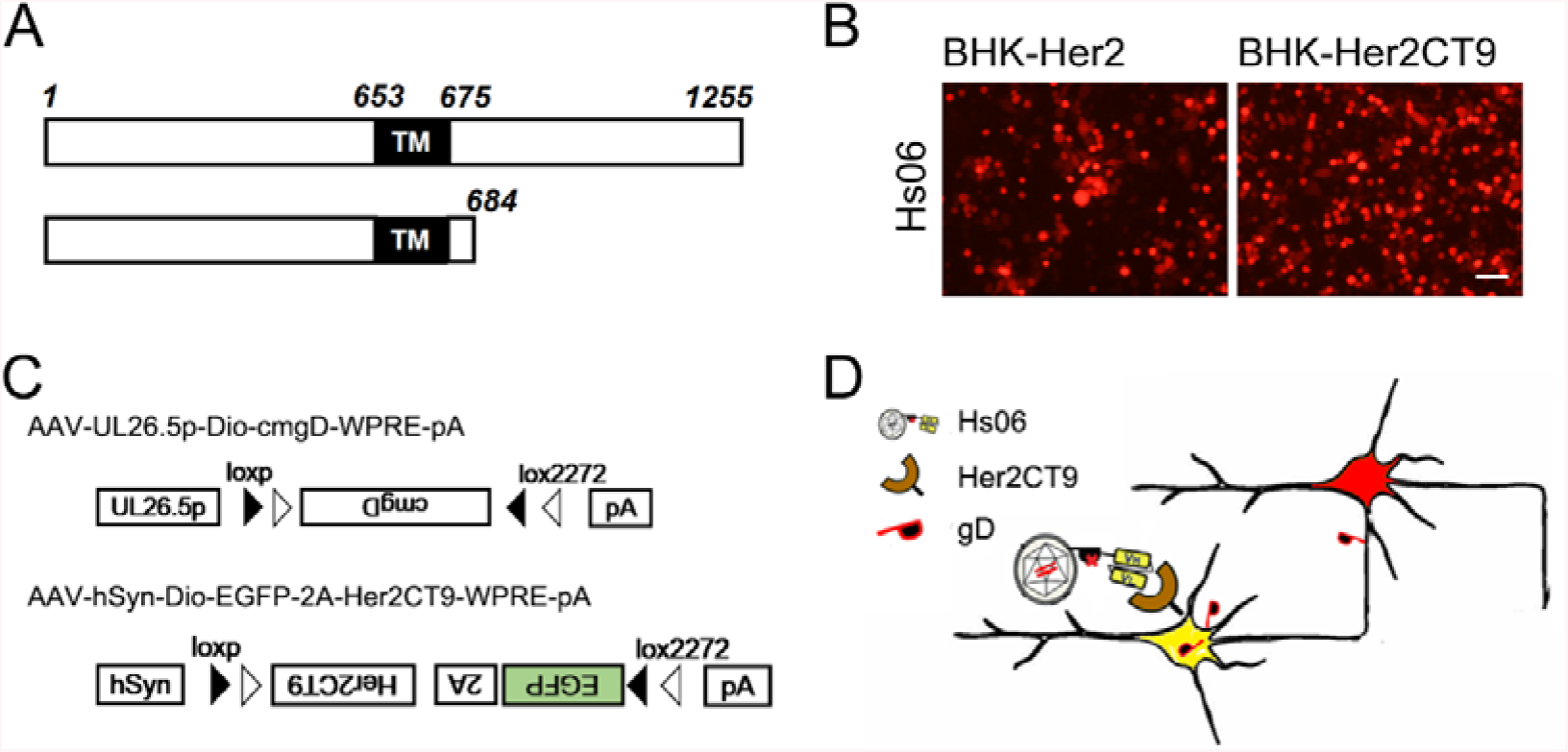
AAV helper construction and strategy of anterograde trans-monosynaptic tracing using Hs06. (A) Structure of Her2 and Her2ct9. TM, transmembrane region. (B) Hs06 recognizes Her2CT9. Hs06 infected BHK-Her2 and BHK-Her2CT9 cells, respectively. Scale bar, 100 μm. (C) Schematic diagram of the AAV helper virus structure required for Hs06 to spread across synapse. To increase the availability of helper virus, Cre-loxp system was used for controlling. (D) Schematic diagram of Hs06 trans-monosynaptic system. The shortened Her2 receptor Her2CT9 and wild-type gD are expressed in neurons in advance with helper viruses. Hs06 recognizes Her2CT9, enters the neuron and then replicates and packages into progeny viruses that could infect the next level of neurons through axon terminals.

### Strategy of anterograde trans-monosynaptic tracing using Hs06

After we realized the specific infection phenotype of H129, then the problem we needed to solve was how Hs06 could infect the postsynaptic neurons after entering the initial neurons. With reference to the rabies virus system, we only needed to compensate the wild-type gD protein in the neurons, then Hs06 could start trans-synaptic spreading. So, a gD expressing AAV should be constructed. We selected the promoter of HSV structural protein gene *UL26.5* (*UL26.5p*) to guide the expression of gD protein, so that only the cells infected by Hs06 would express gD in large quantity ^17^. The use of *UL26.5* promoter could better simulate the protein expression mechanism of wild-type H129 in neurons. In addition, considering that the chimeric protein scHer2/gD in Hs06 contained partial gD sequences, we carried out codon optimization on the *gD* sequence in AAV, and the optimized *gD* sequence was named *cmgD*. In this way, homologous recombination that might exist between Hs06 and the gD-expressing AAV genome was avoided. Finally, the gD-expressing AAV was named AAV-UL26.5p-Dio-cmgD-WPRE-pA (Fig. 2C).

So far, we had completed the design of AAV virus to assist Hs06 virus in specific infection and transmission across synapses. The two viruses were injected into the target brain region in advance, and Hs06 was injected into the same location again after sufficient amounts of Her2CT9 and EGFP were expressed. Hs06 entered into the neurons by recognizing Her2CT9, and then completed the packaging of the progenies with the gD expression element provided by the helper virus, and then infected the postsynaptic neurons through the axon endings (Fig.2D).

### Infection efficiency and specificity of Hs06 in vivo

We had demonstrated that Hs06 did not infect primary cultured neurons in vitro as metioned above. However, it was necessary to verify whether Hs06 was still infection-specific in vivo. So, we injected PBS and the helper virus AAV-hSyn-Dio-Egfp-T2a-Her2CT9-pA into the nucleus accumbens (Nac) brain region of D2R-cre or C57BL/6 mice, respectively (Fig.3A). Two weeks later, Hs06 was injected in situ again, and the brains were collected and sectioned for observation 5 days later. The results showed that a large number of neurons infected by Hs06 could be observed in the Nac and its adjacent areas in the brains of mice which injected Her2CT9 expressing AAV. We did not observe any red neurons in the control group injected with PBS (Fig.3C). In conclusion, the results showed that Hs06 had a high affinity to Her2CT9 and could infect the neurons expressing Her2CT9 efficiently and specifically.

**Figure 3.**
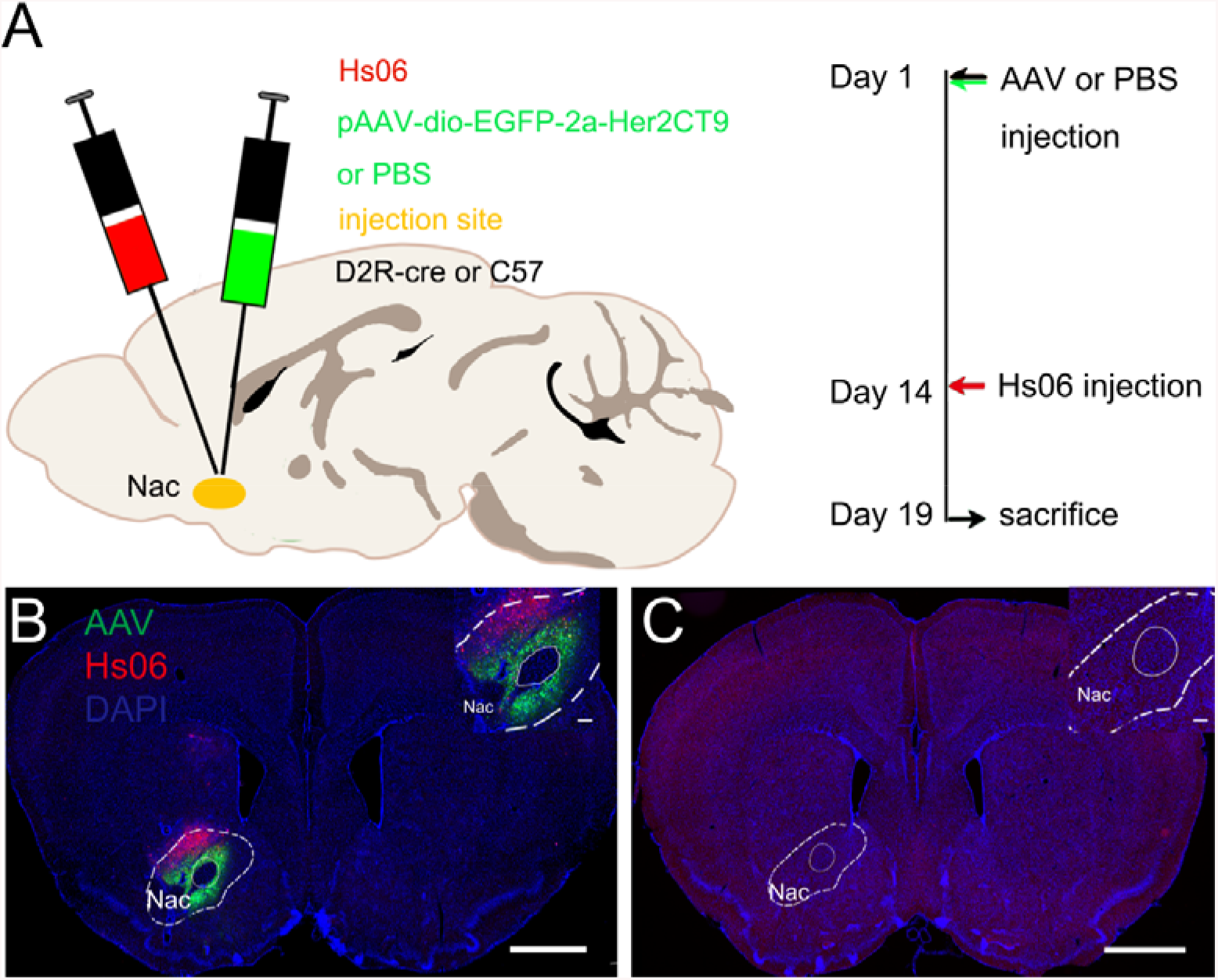
Infection efficiency and specificity of Hs06 in vivo. (A) Schematic diagram of virus injection and experimental procedure. (B) Hs06 could infect neurons that pre-expressed Her2CT9. (C) Hs06 could not infect normal neurons. Scale bar, 1000 μm.

### Hs06 could be used as an anterograde trans-monosynaptic tracer

In aforementioned experiments, we had obtained a theoretically feasible anterograde tracing system combined by Hs06 and its helper virus. Although previous studies had proved that H129 mainly spread anterogradely after entering the cell body of neurons, we still needed to verify whether the new system remained the ability of anterograde transmission due to its modification and recombination.

We performed tracing experiments in the primary visual cortex (V1) of the mouse brain (Fig.4). First, the helper viruses AAV-hSyn-Dio-Egfp-T2a-Her2CT9-pA and AAV-UL26.5p-Dio-cmgD-WPRE-pA were mixed with the Cre-expressing virus AAV-hSyn-Cre-WPRE-pA and injected into the V1 region of C57BL/6 mice. Then, Hs06 was injected into the same site 14 days later. 5 days after Hs06 injection, the brains were collected and sectioned for observation (Fig. 4A). Due to the low fluorescence expression level of Hs06, immunohistochemical treatment was carried out on the brain slices to facilitate imaging and observation. In the labeling results, we were able to observe the expression of the red fluorescent protein in the neurons receiving projections from neurons in V1. These brain regions included the superior colliculus (SC), the lateral geniculate ventral region (LGNv), and the caudal putamen (CPU) (Fig. 4C-F). These brain regions mentioned above had been proved that they only received the projection of V1 but not projected back ^18, 19^, so the neurons labeled in these brain regions should be infected by Hs06 spreading from V1. In the brain regions that were reciprocally connected with V1 (e.g., dorsal lateral geniculate nucleus, LGNd), tdTomato labeling of cell bodies might result from both anterograde and retrograde transport of the virus. These regions were not considered in the current study because of the ambiguity.

**Figure 4.**
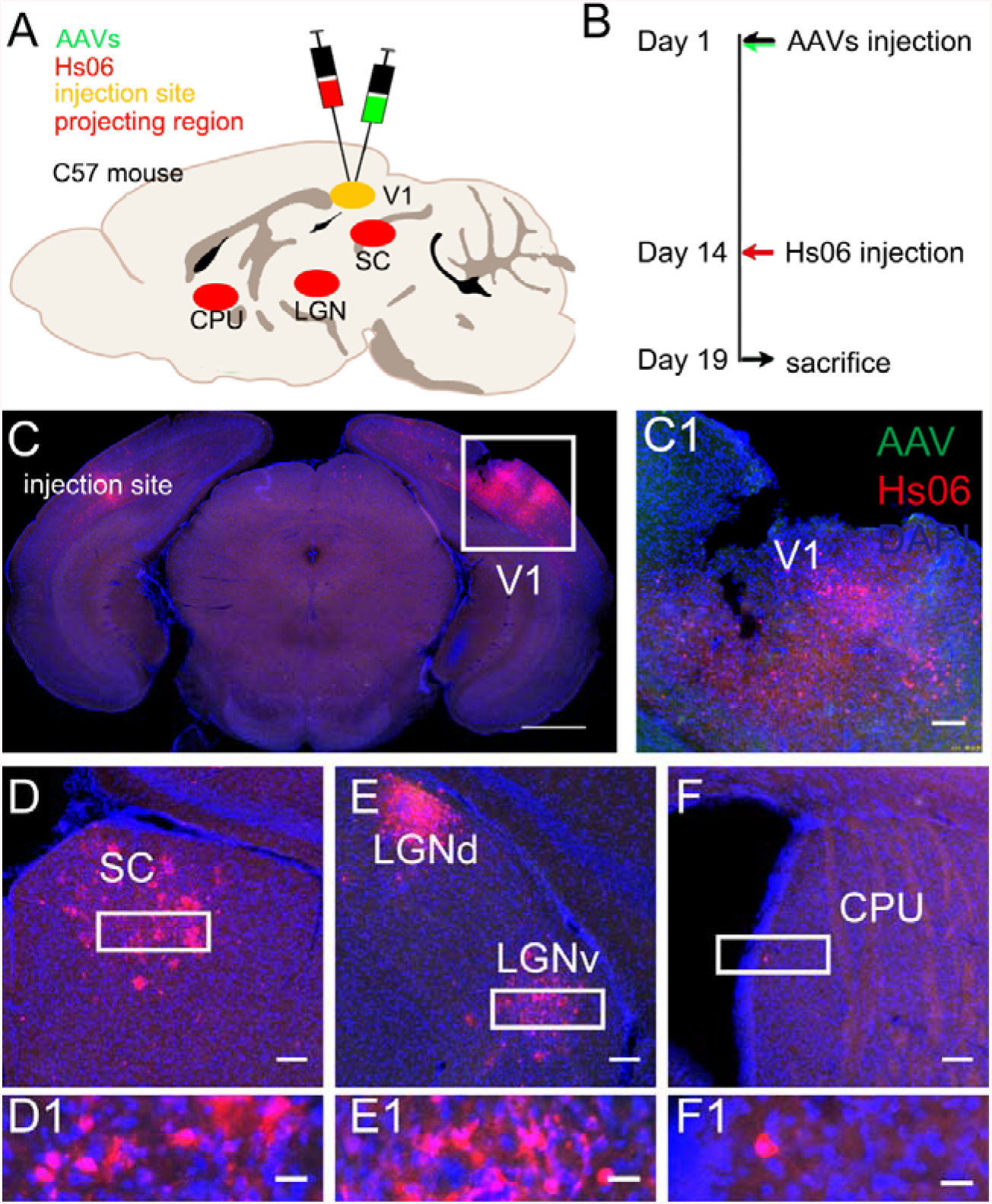
Hs06 could be used as an anterograde trans-monosynaptic tracer. (A) Schematic diagram of main unidirectional projecting brain regions of V1. (B) Time points for virus injection and brain sampling. (C, C1) Image of the injection site V1. Due to the high cytotoxicity of H129, most of the initial neurons in the V1 region have died. (D-F) The downstream brain regions of V1 were labeled by Hs06. Scale bar, C, 1000 μm; D-F, 100 μm; D1-F1, 20 μm.

Furthermore, since Hs06 itself contained chimeric scHer2/gD gene in its genome, whose gene product scHer2/gD protein could also be packaged into complete virions in neurons, so could it transmit to the postsynaptic neurons after entering the neuron without wild-type gD? To answer this question, we mixed AAV-hSyn-Dio-Egfp-T2a-Her2CT9-pA with AAV-hSyn-Cre-WPRE-pA and injected them into V1 to perform the similar experiment as above. The results showed that red fluorescent signals were observed only at the injection site, meaning that Hs06 could enter the cell but not replicate and spread (Fig.S1). In this experiment, due to the lack of wild-type gD, Hs06 could not produce intact infectious virus after entering the neuron, so the green fluorescence signal of the neurons expressing the Her2CT9 receptor could be observed (Fig.S1A).

To further confirm Hs06 virus system could realize anterograde trans-monosynaptic tracing without retrograde spreading, we performed tracing experiment initial form superior colliculus (SC) (Fig.5A). It had been proved that SC only received projection form V1 but not projected back. Neurons in SC project to many regions including pretectal nucleus and LGN ^20^. In this experiment, we observed neurons labeled by Hs06 in SC projecting regions but not V1 (Fig.5C-G). These results suggested that the system could anterograde trans-monosynaptic spread exclusively with the assistance of Her2CT9 and gD. However, because Hs06 was as virulent as the wild type, the initial neurons were mostly dead and difficult to observe at the time of sampling. In addition, we observed very few neurons labeled in CPU, presumably due to the deviation of actual injection site and the trans-synaptic efficiency of the virus system.

**Figure 5.**
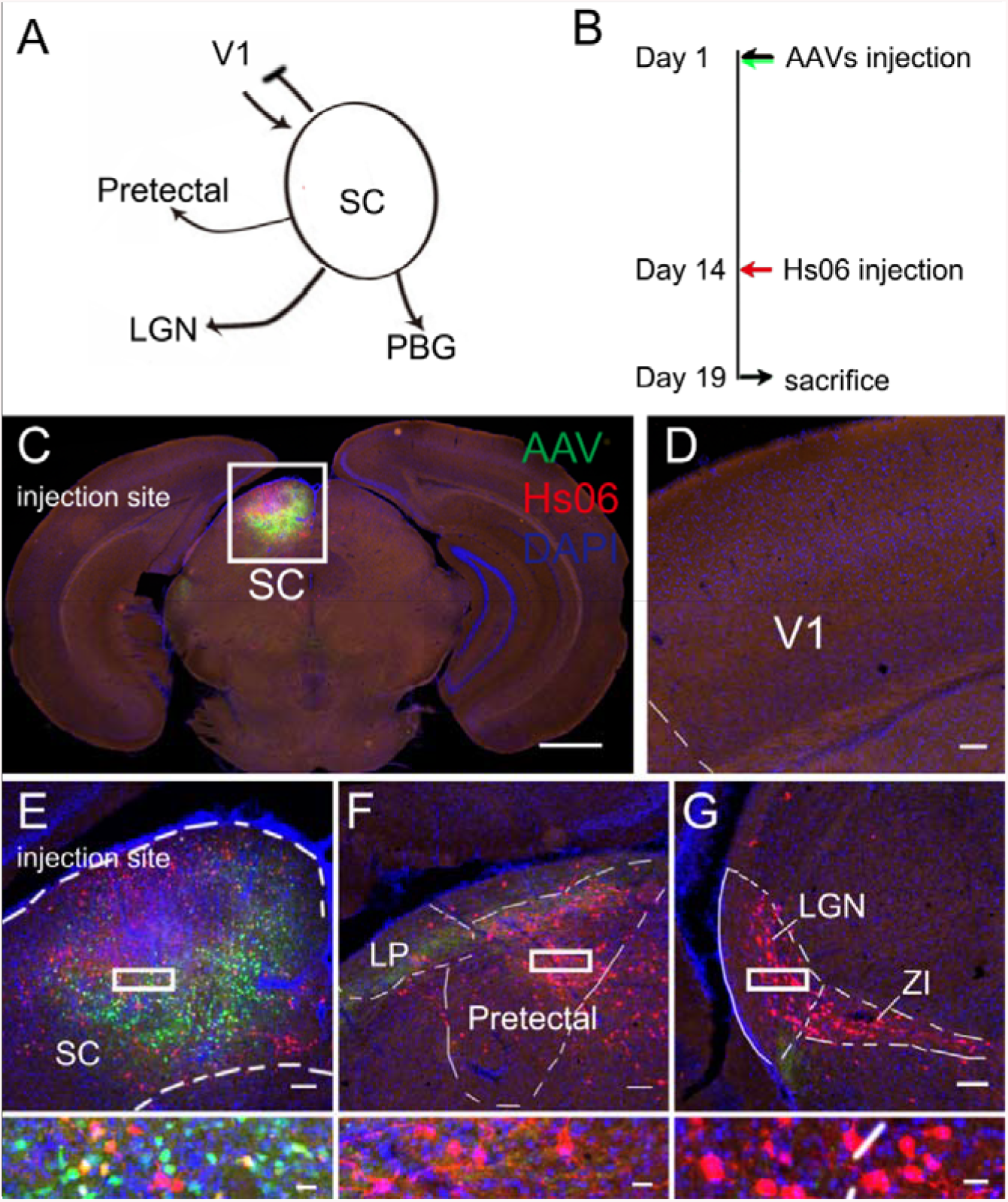
Hs06 could not perform retrograde trans-synaptic spreading. (A) Schematic diagram of main projecting brain regions of SC. SC receives projection form V1 unidirectionally. (B) Time points for virus injection and brain sampling. (C, E) Images of the injection site V1. Due to the high cytotoxicity of H129, few initial neurons in the SC region could be observed at 5 days post-injection. (D) No neurons were labeled by Hs06 in V1. (F, G) The downstream brain regions of SC were labeled by Hs06. Scale bar, C, 1000 μm; D-G, 100 μm; E-G bottom, 20 μm.

### Hs06 dissected the direct output pathway of genetic defined neurons

Precisely mapping the output neuronal circuits required not only the output pathway from given brain regions, but also the projectome information from cell-type specific neurons. We utilized the Hs06 monosynaptic tracing system to dissect the direct output networks of GABAergic neurons in lateral hypothalamus (LH) or ventral tegmental area (VTA).

GABAergic neurons in LH were associated with a variety of instinctive behaviors in animals, such as diet, sleep, and predation ^21^. In recent years, with the development of optogenetics, more and more researches had been conducted on how the GABAergic neurons of LH functions through neural circuits. Nieh et al. activated GABAergic neurons projected from LH to VTA and found that it could drive mice to drink more sugar water ^22^. Cassidy et al. found that GABAergic neurons projected from LH to diagonal band (DBB) could inhibit the neurons of DBB, thus enabling the animals to overcome the anxious environment and cause feeding behavior ^23^. LH was also related to predation and avoidance behavior. Li et al. found that activated LH GABAergic neurons which projected to periaqueductal gray (PAG) could drive mice to prey on moving cockroaches ^24^. The above studies proved that GABAergic neurons of LH could project to multiple brain regions, thus participating in a variety of animal instinctive behaviors. However, these studies mainly used electrophysiology, optogenetics or retrograde tracers to study the connections between neurons. The current labeling methods mainly observed the axonal fiber projection of LH neurons, which were not necessarily the direct synaptic connections. Therefore, we used the Hs06 monosynaptic tracing system to trace the GABAergic neurons of LH (Fig.6). Here, we used GAD2-Cre transgenic mice in tracing experiments. The animal experimental procedure was consistent with the forward tracing experiment of V1, excepted that the helper viruses used here were AAV-hSyn-Dio-Egfp-T2a-Her2CT9-pA and AAV-UL26.5p-Dio-cmgD-WPRE-pA and the injection site was LH (Fig.6A). The labeling results showed that the neurons in the above-mentioned brain regions that received projection from LH GABAergic neurons were labeled in large quantities. Similar with V1 tracing, the initial neurons at the injection site LH basically died, and the initial neurons were not observed (Fig.6B). In other brain regions, we observed the extensive distribution and large number of neurons labeled by Hs06, which indicated that the GABAergic neurons of LH could project to many brain regions, and also proved that Hs06 tracing system had a relatively high trans-monosynaptic spreading efficiency (Fig.6C-E).

**Figure 6.**
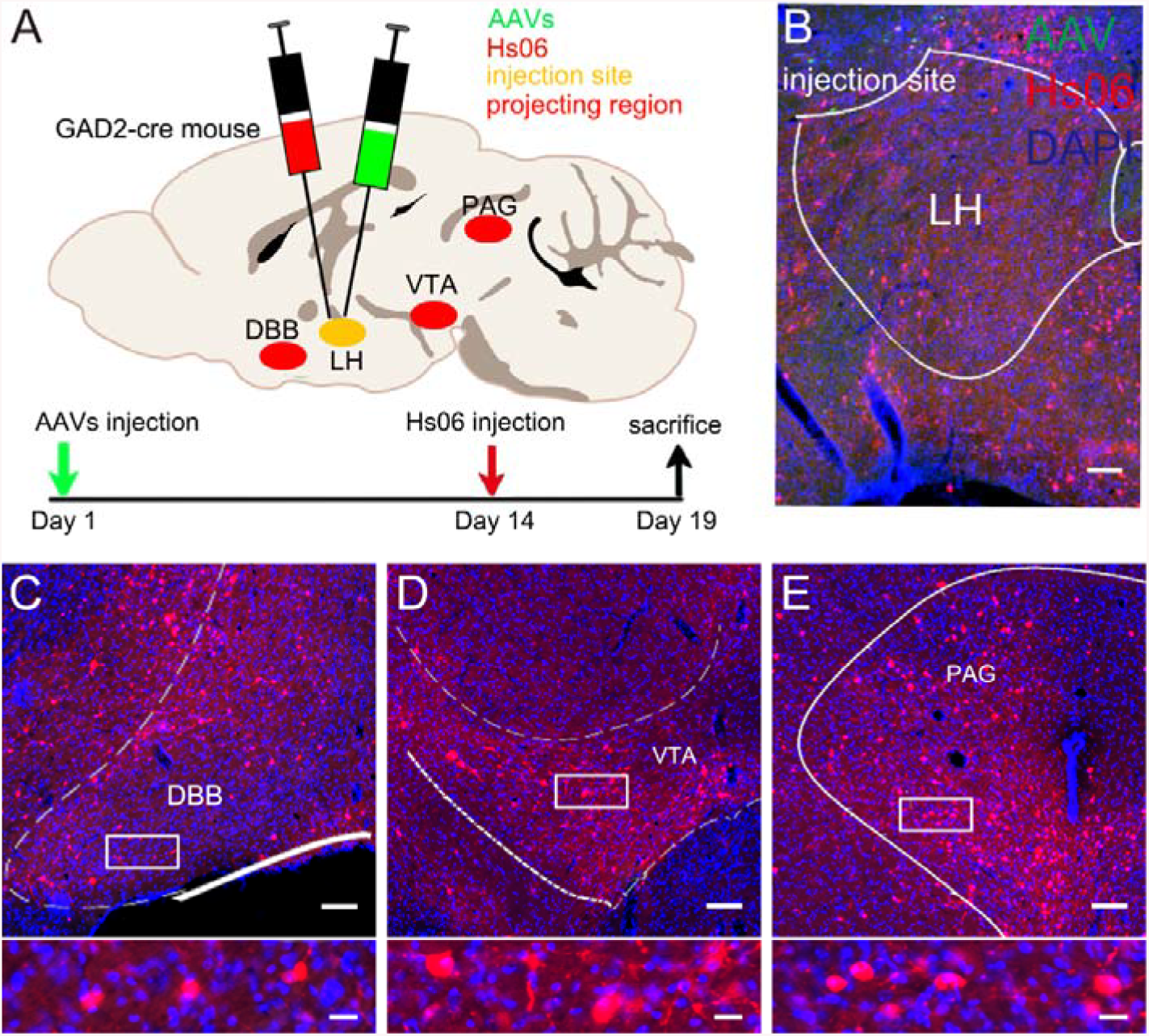
Hs06 dissected the direct output pathway of genetic defined GABA neurons in LH. (A) Main projecting brain regions of LH and the time points for virus injection and brain sampling. (B) The injection site LH. Due to the cytotoxicity of Hs06, the initial neurons were not observed at the time of sampling. (C-E) The downstream brain regions labeled by Hs06. DBB, VTA and PAG were showed respectively. The boxed regions were enlarged to show the neuron morphology. Scale bar, B-E 100 μm; C-E bottom, 20 μm.

In order to further test labeling ability of Hs06 tracing system, we used this system to label the output network of GABAergic neurons of VTA (Fig.7). The GABAergic neurons of VTA could project to many brain regions, such as nucleus accumbens (NAc), ventral globus pallidus (VP), lateral habenula (LHB) and dorsal medial raphe nucleus (DRN), and played an important role in animal behaviors such as pressure and reward ^25^. Applying the tracing system to the VTA of GAD2-Cre transgenic mice, we observed neurons labeled by Hs06 in the above VTA GABAergic neurons projecting brain regions (Fig.7C-E). It was worth noticing that in this experiment, we observed double labeled initial neurons (Fig.7B). This experiment once again verified the anterograde monosynaptic tracing ability of Hs06 system, and also showed that this system had a relatively stable tracing capacity.

**Figure 7.**
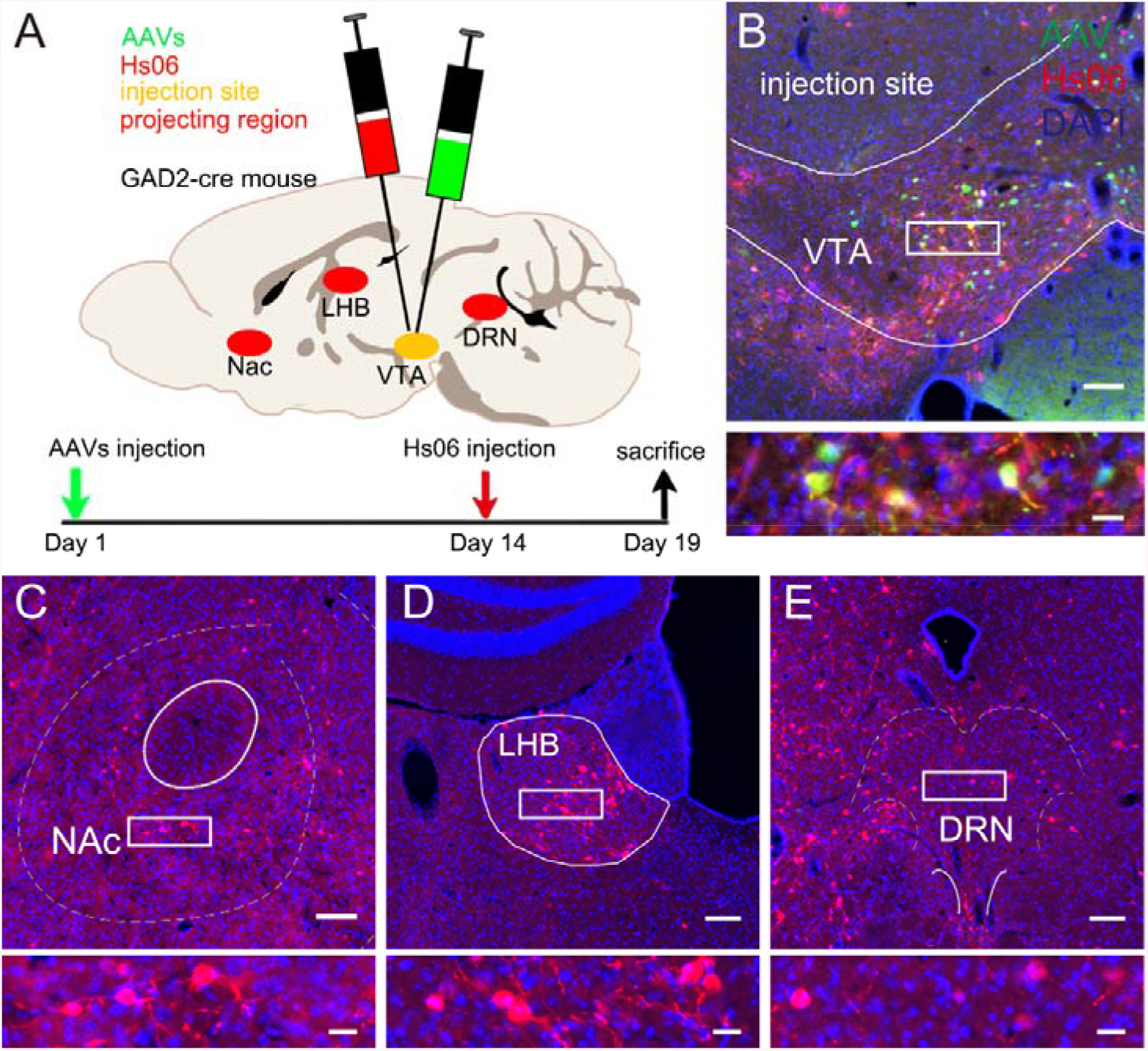
Elucidating the output circuits of VTA GABAergic neurons with Hs06 virus system. (A) Main projecting brain regions of LH and the time points for virus injection and brain sampling. (B) The injection site VTA. Here, double labeled initial neurons were observed. (C-E) The downstream brain regions labeled by Hs06. NAc, LHB and DRN were showed respectively. The boxed regions were enlarged to show the neuron morphology. Scale bar, top, 100 μm; bottom, 20 μm.

The helper AAV could perform slightly retrograde transmission and then expressing Her2CT9 ^26^, thus generating retrograde infection by Hs06, which became a concern. We carried out the control experiment when tracing the output circuits of VTA GABAergic neurons (Fig.S2A). Here we injected Her2CT9 expressing AAV into VTA solely, and then superinfected with Hs06. There were no Hs06 infected neurons observed except the injection site (Fig.S2B-E). The results indicated that the serotype of AAV we used could not conduct effective retrograde transmission to support enough expression of Her2CT9.

## Discussion

Dissecting the brain networks requires appropriate tracing tools, but the anterograde tracing tools are under-development, and particularly, the rigorous monosynaptic anterograde tracer is still lacking ^27^. In this study, we constructed and verified a new monosynaptic tracing system based on the H129 strain of HSV1 ^28^. This system is mainly composed of H129 recombinant Hs06, which can specifically infect neurons expressing the Her2 receptor. Moreover, we tested its function in the known neural circuits, and proved that this system could effectively trace the direct output pathway of genetic defined neurons.

Though H129 had been widely used in anterograde neural circuit tracing ^9, 10, 29, 30, 31, 32, 33, 34^, the axon terminal take-up capacity greatly restricted its further application ^12,35^. Due to this defect, it was hard to start tracing from restricted brain region. So, eliminating axon terminal uptaking of H129 has become an important solution to develop the rigorous monosynaptic anterograde tracer. For this purpose, we were inspired by the strategy used in retargeting of oncolytic HSV (oHSV). Oncolytic HSV is a big field of HSV study and has been developed for decades ^35^. One of the most important research achievements of oHSV is the retargeting strategy ^36, 37^. Researchers introduced single-chain antibody (scFv) that specifically recognize tumor antigens into envelop proteins of HSV to realize tumor targeting and glycoprotein D (gD) was chosen in most cases ^15, 38^. The first-generation retargeted oHSVs carried the retargeting ligand in gD, in place of the receptor binding domain of the glycoprotein. The deleted sequence was critical for gD interaction with the natural viral receptors HVEM (herpesvirus entry mediator) and nectin1. The resulting recombinants were detargeted from these receptors ^36^. The most selected cancer target was human Her2 and the retargeting moiety was a high-affinity scFv ^15, 39^. As Her2 was not expressed in the nervous system, so we chose it as the receptor of retargeting H129 to realize neuron specific infection by providing Her2 artificially. But the complete Her2 receptor gene had a length of 3,768 bp, which was too big to be carried by small capacity AAV vector. By analyzing the protein structure of Her2, we selected to construct the shortened Her2 (Her2CT9), which deleted the long intracellular region of 571 amino acids, remaining extracellular and transmembrane regions of Her2 to adsorb to the cell surface and initiate targeted Hs06 infection process. Using this retargeting strategy, we constructed Hs06, a new H129 recombinant that specially recognized Her2 receptor, and designed as a monosynaptic tracing system.

To simplify the generation and lower the purification difficulty of retargeted H129, we first constructed the gD null H129 strain Hs01 (H129ΔgD-tdTomato). Taking Hs06 as an example, though Hs01 could also be amplified together with Hs06, the progeny viruses were also coated with chimeric protein scHer2/gD expressed by Hs06 genome. So, even if the obtained Hs06 virus contained a small amount of Hs01, the experimental results would not be affected. Therefore, the recombinant virus obtained by recombination with Hs01 did not need to undergo a complex plaque purification process.

It is important to design proper helper vectors for efficient trans-synaptic spreading and tracing. When constructing the gD expressing AAV vector, we performed codon optimization on *gD* gene and selected an unconventional promoter. For the optimization of *gD* codon, we mainly considered to avoid the recombination of *gD* and Hs06 genome to form a wild type virus. The reason was that the chimeric protein scHer2/gD was surrounded by two fragments of gD ^14^. Therefore, Hs06 and wild-type *gD* have a certain chance of recombination. Codon optimization of *gD* could avoid this possibility. For the choice of promoter expressing *gD*, we considered to better simulate the virus replication of wild-type H129. Since *gD* is a late gene of HSV virus, we selected the promoter of *UL26.5*, which was also a late gene, to guide its expression.

With the Cre transgenic mouse lines, the Hs06 anterograde monosynaptic tracing system offers a potential tool to dissect the direct projectome of a specific type of neuron in a given brain region. Before mapping the direct projectome using Hs06 and the helpers, it was necessary to test the infection specificity. In addition, it had been reported that AAV also had the ability to transport retrogradely. Therefore, in the tracing experiments without providing gD, we observed the brain regions projected to VTA, such as the lateral habenula (LHB), and found no red fluorescence signal, thus excluding the retrograde infection of Hs06 caused by the reverse absorption of AAV (Fig.S2). These controlled experiments showed that the Hs06 system spreaded across synapses only when gD expressing helper virus was provided and no longer had the reverse infectivity of wildtype H129.

Though Hs06 monosynaptic tracing system can be used as an anterograde tracer, there are still defects. The biggest one is that the starter neurons can hardly be observed when using this system. Because we only modified the infection characteristics of H129 while without reducing its toxicity, Hs06 was still as cytotoxic as its prototype virus strain H129. Thus, the initially infected neurons were killed and cleaned, which led to difficulty in observing of starter neurons. Can we observe the starter neurons if we shorten the sampling time? Indeed, after shortening the sampling time from 5 days to 3 days, we observed the signal of the initial neurons, but the trans-synaptic efficiency was reduced and the fluorescence signal of the post-synaptic neurons was faint (data not shown). The follow-up optimization work in virus attenuation of H129 and enhancement of exogenous gene expression will further improve the trans-monosynaptic tracing efficiency of Hs06 system.

In summary, we have developed a rigorous anterograde trans-monosynaptic tracer derived from HSV1 H129 strain. Hs06 represents a potential novel anterograde monosynaptic tracer, which may contribute in revealing the direct projectome connectivity. This tracer complements the current neuronal circuit tracer tool box.

## Methods

### Animals

All husbandry and experimental procedures in this study were approved by the Animal Care and Use Committees at the Shenzhen Institute of Advanced Technology (SIAT) or Wuhan Institute of Physics and Mathematics (WIPM), Chinese Academy of Sciences (CAS). Adult male C57BL/6 mice were purchased from Hunan SJA Laboratory Animal Company. The GAD2-cre mice used were heterozygote and generated by mating the transgenic male mice with C57BL/6 female mice. All animals were housed in a dedicated housing room with a 12/12 h light/dark cycle, and food and water were available ad libitum. All the experiments with viruses were performed in bio-safety level 2 (BSL-2) laboratory and animal facilities.

### Construction of gD and Her2 expressing cell lines

Firstly, we constructed a lentiviral vector containing the *gD* gene, the *gD* gene was amplified from H129 genome using specific primers. The PCR product was inserted into FUGW (addgene, #14883) to create pFUGW-HisEgfp-T2A-gD. Then, the plasmid pFUGW-HisEgfp-T2A-Her2 was constructed by replacing the *gD* gene with the *Her2* gene. The *Her2* gene was amplified from plasmid pBABEpuro-ERBB2 (addgene, #40978) using specific primers. Lentivirus harboring the *gD* or *Her2* gene was generated based on the method described previously ^40^ and used to transduce BHK-21 cells. In brief, the above plasmids were co-transfected with three lentivirus packaging vectors (pGAG/POL, pREV and pMD2.G) into 293T cells, and the targeting viruses were obtained by collecting supernatants after 2 to 3 days. The supernatants were used to transduce well cultured BHK-21 cells in 6-well plate respectively, and the proportion of cells expressing nuclear located green fluorescence was observed two days later. Cells in the well which most of the cells expressing fluorescence were taken for subculture, and the cell lines obtained were named BHK-gD and BHK-Her2, respectively.

### Construction of gD null H129 strain

To obtain the *gD* null H129 strain, termed Hs01, we first constructed a *gD* targeting shuttle vector. Using H129 genome as templates we amplified the upstream and downstream sequence of *gD* gene to be the homologous arms. Then the two PCR products were inserted into pcDNA3.1^+^ together to get plasmid pcDNA3.1^+^-ΔgD. And then red fluorescent protein tdTomato expressing box hUbc-tdTomato-WPRE-pA was inserted between the two homologous arms to construct the *gD* targeting shuttle vector pcDNA3.1^+^-ΔgD-hUbc-tdTomato-WPRE-pA, which was named pHs01. To generate the recombinant Hs01, pHs01 was purified by Omega plasmid mini kit and co-transfected with H129 genomic DNA into 293T cells in six-well plates, using Lipofectamine 2000 (Invitrogen) according to the manufacturer’s instructions. After the majority of cells showed cytopathic effect, the medium was removed and the cells were harvested in PBS. After three-rounds of freeze-thaw-vortex, the cell lysate was used to infect BHK-gD cells plated in six-well plates. After 1 h of infection, viruses were removed and DMEM with 2% FBS, antibiotics, and 1% agarose were overlaid on the cells. After 2 or 3 days, well separated EGFP-expressing plaques were picked and subjected to at least five more rounds of plaque purification to remove wild type H129 virus.

### Construction and production of Her2 retargeting H129 recombinant

In order to generate the Her2-retargeting H129 recombinant, termed Hs06, the first step was to construct a Her2-targeting *gD* gene. To construct the Her2-targeting *gD* gene, we need to select a single chain antibody (scHer2) with a high affinity for HER2. In this paper, we selected the *scHer2* sequence reported by Xiaodan Cao et al. and uploaded to NCBI (GenBank: KM016462.1) ^41^. The chimeric protein scHer2/gD was formed by using this sequence to replace the 6-38 amino acids of the gD protein, the key part of the gD protein that recognizes the natural receptor for HSV. Subsequently, we constructed the shuttle vector phUbc-mCherry-t2a-scHer2/gD-WPRE-pA which named pHs06. Then, homologous recombination was performed to generate Hs06 via transfecting shuttle vector into BHK-Her2 cells and superinfecting with Hs01. Here, the upstream and downstream homologous arms used for recombination were the hUbc and WPRE sequences, respectively. Hs06 was purified using plaque assay described above.

Purified Hs06 viruses were mass-produced by infecting BHK-her2 cells grown in 10 cm plates (M.O.I = 0.1~0.01). After infected cells showed a prominent cytopathic effect (~3 days), medium containing the viruses was collected from about twenty 10 cm plates, spun down to remove cell debris (6,000 rpm for 5 minutes), the supernatant passed through a 0.22 μm filter, and finally centrifuged at 25,000 rpm/1.5 hours in a JA25.50 rotor using a Model L80XP Beckman Coulter Ultracentrifuge. The virus pellet was resuspended overnight at 4 °C in a small amount of cold PBS (PH=7.4) (~0.2 ml) with constant shaking. Dissolved viruses were aliquoted into 3 μl and stored in 200 μl PCR tubes at −80 °C. The titer of viral stocks was determined using standard plaque assay on BHK-Her2 cells and titers were expressed as plaque-forming units (PFU) per milliliter. A fresh aliquot of stock virus was thawed and used for each experiment. The titer of Hs06 used in these studies was ~1 × 10^8^ PFU/ml.

### Preparation of AAV helpers

First, we purchased the AAV skeleton plasmid containing the Cre-Loxp system from addgene (pAAV-hSyn-Dio-mCherry-WPRE-pA, #44361). Based on this plasmid, we modified and inserted the required expression elements into it. To construct the plasmid pAAV-hSyn-Dio-Egfp-T2a-Her2ct9-pA, we inserted *Egfp-T2a-Her2ct9* into the Cre-Loxp element to replace the original mCherry and then deleted WPRE. To construct pAAV-UL26.5p-Dio-cmgD-WPRE-pA, we first replaced *mCherry* with *cmgD*, and then replaced the original hSyn promoter with *UL26.5* promoter.

AAV was prepared by transfection HEK293T with the three plasmids system as described before ^42, 43, 44^. HEK293T was maintained in DMEM medium with 10% FBS, 1% penicillin and 1% streptomycin, and cultured in an incubator at 37 L and 5% CO_2_. HEK293T cells were cultured in 15 10-cm dishes 24 hours before transfection, and DMEM medium containing 2% FBS was used to culture the cells until the confluence was about 80% at the time of transfection. All plasmids and PEI transfection reagent were added to serum-free DMEM medium in a 1.5 ml EP tube. The ratio of plasmid DNA to transfection reagent was 1:2. After mixing, the plasmid DNA was left at room temperature for 15-20 minutes, and then added to the dish drop by drop.72 hours after transfection, supernatants and cells were collected, respectively. The collected HEK293T cells containing AAV were precipitated and re-suspended to 1×10^7^ cells/mL with cell lysate. After repeated freezing and thawing in liquid nitrogen and water bath for 3 times, an appropriate amount of nuclease Benzonase (1 μL/10 mL) was added, fully mixed, and digested at 37 ◻ for 1 hour. The supernatant was then collected by centrifugation at 2500 g for 10 minutes. The supernatant of lysate treated by nuclease and the supernatant of cell culture after transfection was mixed together with 8% PEG8000 and 0.5 mol/L NaCl solution, then placed at 4 L overnight, and centrifuged at 4 L at 12000 g for 90 minutes. Blown and dissolved the precipitation in 10mL PBS to obtain the initial concentration of the virus. Then, iodixanol density gradient centrifugation was used to concentrate secondarily. The final virus concentrate was filtered and sterilized with a 0.22 micron filter, then stored in at −80 ◻. QPCR was used to q the titer of AAV, and the skeleton plasmid was diluted in 10 fold gradient to make the standard curve.

The titer of AAVs used in this study were 1.3 × 10^13^ GC/mL for AAV-hSyn-Dio-EGFP-T2a-Her2CT9-pA, 1 × 10^13^ GC/mL for AAV-UL26.5p-Dio-cmgD-WPRE-pA and 2.8 × 10^12^ GC/mL for AAV-hSyn-Cre-WPRE-pA.

### Mice and viral injections

All procedures on animals were performed in Biosafety level 2 (BSL-2) animal facilities as before ^45, 46^. Animals were anesthetized with pentobarbital sodium (80 mg/kg, i.p.), and placed in a stereotaxic apparatus (Item: 68030, RWD, Shenzhen, China). The skull above the targeted areas was thinned with a dental drill and removed carefully. Injections were conducted with a syringe pump (Item: 53311, Quintessential stereotaxic injector, Stoelting, United States) connected to a glass micropipette with a tip diameter of 10-15 mm. The glass micropipette was held for an extra 10 min after the completion of the injection and then slowly retreated. After the surgery, the incisions were stitched and lincomycin hydrochloride and lidocaine hydrochloride gel was applied to prevent inflammation and alleviate pain for the animals.

For V1 injection, the mixture of AAV-ul26.5p-Dio-cmgD-WPRE-pA, AAV-hSyn-Dio-Egfp-T2a-Her2ct9-pA and AAV-hSyn-Cre-WPRE-pA (volume ratio: 8:2:1, 100 nl in total) was injected into the adult C57 mice with the following coordinates: AP, −3.88 mm; ML, −2.60 mm; and DV, 0.5 mm ventral from the cortical surface. Then, 200 nl Hs06 was injected into the same injection site with the AAV mixture after 2 weeks. Five days after the Hs06 injection, the mice were perfused for brain collection.

For LH and VTA injection, the mixture of AAV-ul26.5p-Dio-cmgD-WPRE-pA and AAV-hSyn-Dio-Egfp-T2a-Her2ct9-pA (volume ratio: 4:1, 100nl in total) was injected into the adult GAD2-cre mice with the following coordinates: AP, −0.82 mm; ML, −1.20 mm; DV, −5.00 mm for LH and AP, −3.20 mm; ML, −0.25 mm; DV, −4.40 mm for VTA. Then, 200 nl Hs06 was injected into the same injection site with the AAV mixture after 2 weeks. Five days after the Hs06 injection, the mice were perfused for brain collection.

### Slice preparation and immunohistochemistry

The mice were anesthetized with pentobarbital sodium (50 mg/kg body weight, i.p.), and perfused transcardially with PBS (5 min), followed by ice-cold 4% paraformaldehyde (PFA, 158127 MSDS, sigma) dissolved in PBS (5 min). The brain tissues were carefully removed and post-fixed in PBS containing 4% PFA at 4 °C overnight, and then equilibrated in PBS containing 25% sucrose at 4 °C for 72 h. The 40 mm thick coronal slices of the whole brain were obtained using the cryostat microtome and stored at −20 °C.

For immumohistochemical staining, sections were rehydrated with PBS, blocked with 10% goat serum in PBS containing 0.1% Triton-X100 (PBST), followed by overnight incubation with primary antibody diluted in PBST. The primary antibody used in this study were goat anti-DsRed (Takara, 1:1000). Sections were washed with PBS and reincubated with CY3-conjugated secondary antibodies (1:200 dilution, sigma). One hour later, sections were stained with DAPI, washed with PBS, mounted with 70% glycerol (in PBS) and sealed with nail polish. For all samples, every sixth section of the brain slices were selected. All of the images were captured with the Olympus VS120 virtual microscopy slide scanning system (Olympus, Shanghai, China).

## Acknowledgements

We thank Professor Lynn Enquist (Princeton University, Princeton, NJ) and Professor David J. Anderson (California Institute of Technology, Pasadena, CA) for providing the HSV-1 H129 viral strains. This work was supported by National Natural Science Foundation of China (31771198, 91632303 and 91732304), Science and Technology Planning Project of Guangdong Province (2018B030331001), CAS Key Laboratory of Brain Connectome and Manipulation (2019DP173024), Guangdong Provincial Key Laboratory of Brain Connectome and Behavior (2017B030301017), and The Strategic Priority Research Program of Chinese Academy of Sciences (XDB32030200).

## Author contributions

PS, HW and FX generated the idea, PS, MY, HW, JX,YW and YL performed the experiments, PS and HW analyzed the data, PS, HW and FX conceived the manuscript and wrote the text, and PS generated the figures.

