## Supplemental Figures and Table for "Rigorous anterograde trans-monosynaptic tracing of genetic defined neurons with retargeted HSV1 H129"

**Supplemental information**

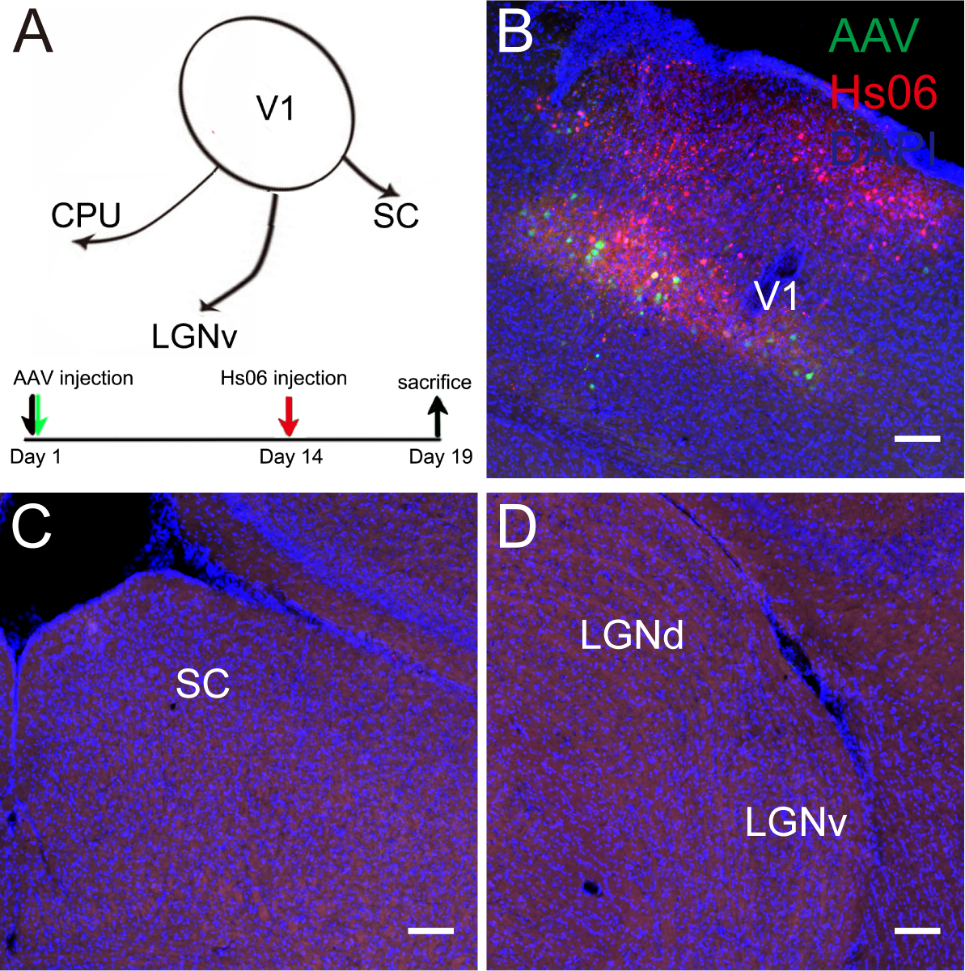

Figure S1. No trans-synaptic transmission occured after Hs06 entered neurons alone. (A) The injection site V1. Because gD was not provided, Hs06 could not undergo lytic infection, so the green fluorescent signal of the neurons expressing the Her2CT9 receptor could be observed here (B). (C, D) No red fluorescence signal was observed in the V1 projection regions SC and LGN. Scale bar, 100 µm.

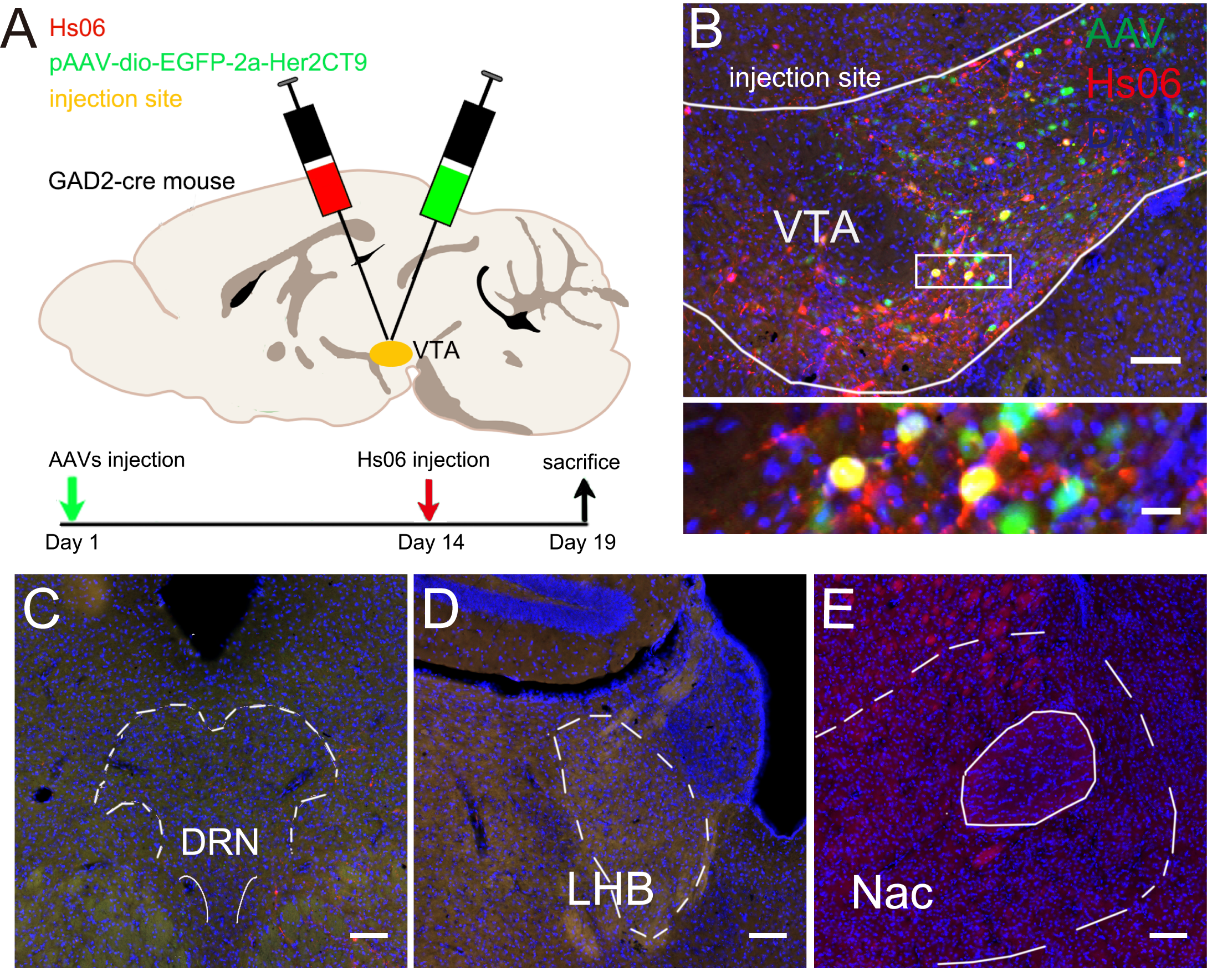

Figure S2. The serotype of AAV we used did not exhibit retrograde transmission. (A) Main projecting brain regions of LH and the time points for virus injection and brain sampling. (B) The injection site VTA. (C-E) The downstream brain regions labeled by Hs06. NAc, LHB and DRN were showed respectively. Scale bar, B-E 100 µm; B bottom, 20 µm.

Table S1 Summary of viral tools used in this study.

| Virus | Serotype | titer |
| --- | --- | --- |
| AAV-Hsyn-DIO-EGFP-2A-Her2CT9-pA | AAV9 | 1.3E+13 GC/mL |
| AAV-UL26.5p-DIO-cmgD-WPRE-pA | AAV9 | 1.0E+13 GC/mL |
| AAV-hSyn-Cre-WPRE-pA | AAV9 | 2.8E+12 GC/mL |
| Hs06 | HSV1-H129 | 1.0E+8 PFU/mL |
| Hs01 | HSV1-H129 | 1.0E+6 PFU/mL |

The viral strain, serotype and titer of virus vectors used in this study. AAV, adeno-associated virus; HSV, herpes simplex virus. GC, genome copy; PFU, plaque forming unit.
